# Exploring nematode diversity in soils: comparison of extraction methods and sequencing protocols

**DOI:** 10.64898/2026.09.29.755246

**Authors:** Marco Fioratti Junod, Sabine Brodbeck, Sophie Gombeer, Lisa van Sluijs, Joost A.G. Riksen, Beat Frey, Anne Kempel, Christian Rixen, Marcus Schaub, Anita C. Risch, Irene Cordero

**Author notes:** Corresponding author:* Irene Cordero Swiss Federal Institute for Forest, Snow and Landscape Research WSL Zürcherstrasse 111, CH-8903 Birmensdorf, Switzerland.

## Abstract

Soil nematodes play key roles in regulating organic matter decomposition, nutrient cycling and trophic interactions in the soil ecosystem, and are very useful bioindicators of soil condition and soil health. But to characterise them, we need reliable measurements of nematode biodiversity in soils. DNA-based techniques such as metabarcoding have recently explored their biodiversity, with successful results. However, there are many methodological steps that can modify the outcome considerably and require informed decisions. In this study, we tested the effect on nematode alpha and beta diversity metrics of three methodological steps, namely i) type of extraction of genetic material from the matrix, either from soil directly with different starting amount of material or extracting nematodes first via the Baermann or Oostenbrink methods, ii) the choice of taxonomically-relevant DNA segment for amplification, with a nematode specific primer (NemF/18Sr2b) and a universal eukaryotic primer (3NDf/1132rmod), both within 18S rDNA, and iii) the selection of appropriate reference library for taxonomic assignment (comparing three nematode specialised databases and three broad taxonomic databases).

Observed differences in alpha diversity were more pronounced when comparing the nematode extraction *vs* soil extractions. In particular, direct soil extractions co-extracted many non-nematode taxa, especially if amplified with a universal primer, which can significantly reduce the number of nematode reads obtained and therefore the accuracy of nematode community indexes and other nematode-based metrics. The choice of the reference database strongly affected the taxonomic identification, which calls for extreme caution when making this selection. Notably, specialised databases tended to over assign reads to nematodes that were detected as non-nematodes by broad taxonomic databases. However, beta diversity metrics were consistent across methods, highlighting these metrics as robust against methodological choices.

Our study clearly demonstrates that the choices of methods for nematode biodiversity studies using molecular tools can have a big impact on the results, and decisions should be made consciously and carefully.

## Introduction

Nematodes are considered the most abundant animal phylum on our planet, with an estimated abundance of 440 quintillion individuals in the first 15 centimetres of soil (van den Hoogen et al., 2019, 2020). The group is characterised by great ecological, functional and morphological variety (Lazarova et al., 2021), responds to edaphic, latitude and altitude gradients (Nisa et al., 2021) and promptly reconfigures in the face of land-use and climate change (Pires et al., 2023). Their role is pivotal in ecosystem services worldwide (Trap et al., 2025), including nutrient cycling (Ghaderi et al., 2026), agricultural production (Na et al., 2025) and pathogen control (Y. Zhang et al., 2020). This makes them perfect candidates as ecological indicators in natural and agricultural soils (Gao et al., 2020; J. Liu et al., 2022; L. Liu et al., 2021; Lu et al., 2020) and a variety of indices have been developed based on the structure and composition of their communities (Biswal, 2022; Du Preez et al., 2022; C. Zhang et al., 2024).

The traditional, morphology-based techniques to characterise nematode communities are increasingly difficult to implement due to a shortage of highly specialised taxonomical skills and the time intensiveness of these processes which preclude high-throughput applications (Geisen et al., 2018; Griffiths et al., 2018). Amplicon-based molecular techniques such as metabarcoding promise deep taxonomic recovery and highly scalable results at a fraction of the cost of microscopy-based expert characterisation (Ficetola et al., 2024; Hayden et al., 2025). However, environmental metabarcoding pipelines involve a long series of steps and choices with important consequences; including extraction of genetic material from the matrix (Waeyenberge et al., 2019), the choice of taxonomically-relevant DNA segment for amplification (Akanwari et al., 2026), bioinformatic filters (Alberdi et al., 2018) and the selection of reference library for taxonomic assignment (De Santiago et al., 2025). All stages have been shown to have consequences for the quantification of richness and diversity indices and detection of rare taxa, as well as the detection of ecological signals which is often neglected in methodological studies.

Soil nematode molecular pipelines differ first of all in the approach used to separate nematode DNA from the matrix. Direct extraction from soil is reported to achieve good depth and representation of nematode soil communities in combination with appropriate primers and protocol (Akanwari et al., 2026; Kenmotsu et al., 2021; Sapkota & Nicolaisen, 2015). However, competitive amplification of non-target sequences and the presence of inhibiting compounds in the soil are reported to be capable of compromising results (Donhauser et al., 2023; Omer et al., 2022). The quantity of soil used for the extraction appears to be critical, with studies reporting species richness to be correlated with the amount of soil processed (Kageyama & Toju, 2022). Isolating nematodes prior to DNA extraction to increase the proportion of target DNA is also performed with a variety of methods. Active methods are based on the facilitation of the spontaneous migration of nematodes from soil to a collecting vessel. The most widespread of these techniques, also in a metabarcoding context, is the Baermann funnel (Baermann, 1917), whereby the soil is laid in a funnel, on a porous membrane, and a humidity gradient is created by regularly wetting the soil. Active nematodes move into the water and sink to the bottom of the funnel. Nematodes are collected after a variable period of time (usually between 24 and 96 h) by unclamping the stem and draining the liquid into a receptacle. Only individuals that are alive, active and capable of movement are therefore collected. On the other hand, passive methods involve separating the nematodes from the soil matrix by exploiting differences in specific weight. Sucrose flotation (Akanwari et al., 2026) is occasionally used in metabarcoding pipelines, but to avoid inhibiting possible amplification inhibiting effects of flotation solutions other than water, elutriators are preferred. Elutriators are hydraulic contraptions generating a tightly controlled upward flow of water on the precipitating stream of soil that allows nematodes to be differentially suspended from soil particles and separately collected, prior to sieving and concentration. The Oostenbrink elutriator (Greco & Crozzoli, 2024; Tintori et al., 2022) is the most commercially successful and widely adopted of these implementations and the best candidate for standardisation. This type of elutriation is purportedly able to separate also inactive or slow-moving individuals. Fine tuning of both the

Oostenbrink elutriator (Verschoor & Goede, 2000) and the Baermann funnel (Cesarz et al., 2019) methods can significantly improve the extraction efficiency.

After DNA extraction, the choice of DNA fragment used for amplification also comes with consequences. The main trade-off is between universal eukaryotic or metazoan primers and targeted-nematode primers. The most advanced versions of the former capture a broader range of organisms (Geisen et al., 2018; Kenmotsu et al., 2021), which may be relevant to the analysis and provide useful ecological insights, while providing sufficient coverage for nematodes. However, the volume of non-nematode sequences can drown the signal of target sequences, and even other metazoan phyla may be better characterised with more specific primers (Akanwari et al., 2026; Sun et al., 2023). Within both categories, while COI has been proposed as a possible alternative (Ren et al., 2023), the 18S and ITS rDNA regions are usually favoured for metabarcoding applications concerning nematodes. State-of-the-art specific primers have achieved 74% of nematode reads (Sikder et al., 2020) and represent a potentially substantial step forward compared to older standards (Porazinska et al., 2009).

The taxonomic resolution of sequences based on comparison with reference libraries is also a non-neutral undertake. It has been suggested that universal or broad taxonomic libraries often lack the volume and specificity to cover nematodes (Ahmed et al., 2019; Baker et al., 2023; De Santiago et al., 2025). For this reason, a series of specialised tools comprising databases and assignment bioinformatic pipelines have been developed (Gamble et al., 2021; Gattoni et al., 2023; Workentine et al., 2020). However, the performance of these novel tools has not been thoroughly assessed in a comparative way.

Even more relevantly, although extraction methods, primer bias, and reference-library completeness are individually recognized as important sources of variation, they are most often evaluated separately and not as part of complete pipelines. It is reasonable to expect strong interaction effects across the sequential steps in the process, with target organism proportions in the extracted DNA affecting amplification performance, and in turn sampled community make-up affecting the taxonomic resolution offered by the different libraries for target and non-target phyla. This can have radical consequences for both the taxonomic description of soil nematode communities and the recovery of experimentally imposed environmental differences. Our study has the ambition to address this problem by comparing together: a) five extraction methods (Oostenbrink extraction plus Qiagen DNeasy Blood and Tissue kit, Baermann extraction plus Qiagen DNeasy Blood and Tissue kit, bulk soil plus small volume Qiagen DNeasy PowerSoil Pro, bulk soil plus small volume Qiagen DNeasy PowerSoil Pro with an intermediate buffered slurry stage, bulk soil plus large volume Qiagen DNeasy PowerMax Soil); b) two state-of-the-art primer pairs – one targeted at nematodes (NemF/18Sr2B, Sapkota & Nicolaisen, 2015; Sikder et al., 2020) and the other at metazoans at large (3NDf/1132rmod, Geisen et al., 2018); c) six reference libraries and assignment protocols, three broadly eukaryotic (Silva132, Quast et al., 2013; Eukaryome, Tedersoo et al., 2024; PR2, Guillou et al., 2013) and three nematode-specific (NemaBase, Gattoni et al., 2023; Nemabiome, (Workentine et al., 2020; NemaTaxa, Baker et al., 2023). The whole pipelines were tested on soil samples from three experimental sites across Switzerland, each featuring a control and an enrichment treatment, in order to test the potential to detect authentic ecological signals.

The questions that our study is planned to give an answer to are:

i. How do nematode-concentration and direct-soil DNA-extraction methods differ in specimen recovery and in the richness, diversity, and composition detected by metabarcoding?
ii. How does primer specificity influence nematode detection, and does its effect depend on the DNA-extraction method?
iii. Which extraction–primer combinations most consistently recover the community response to enrichment across contrasting sites?
iv. How do generalist and nematode-specialized reference libraries differ in taxonomic coverage, resolution, composition, and accuracy?

## Material and methods

### Experimental sites and soil collection

The soil samples were collected from three long-term experimental sites in Switzerland (Figure 1a). In all cases, the experimental design is more complex, but the sampling was limited to a subset of plots representing enriched (with irrigation or fertiliser application) and control conditions. For each of the sites three enriched and three control plots were chosen.

**Figure 1:**
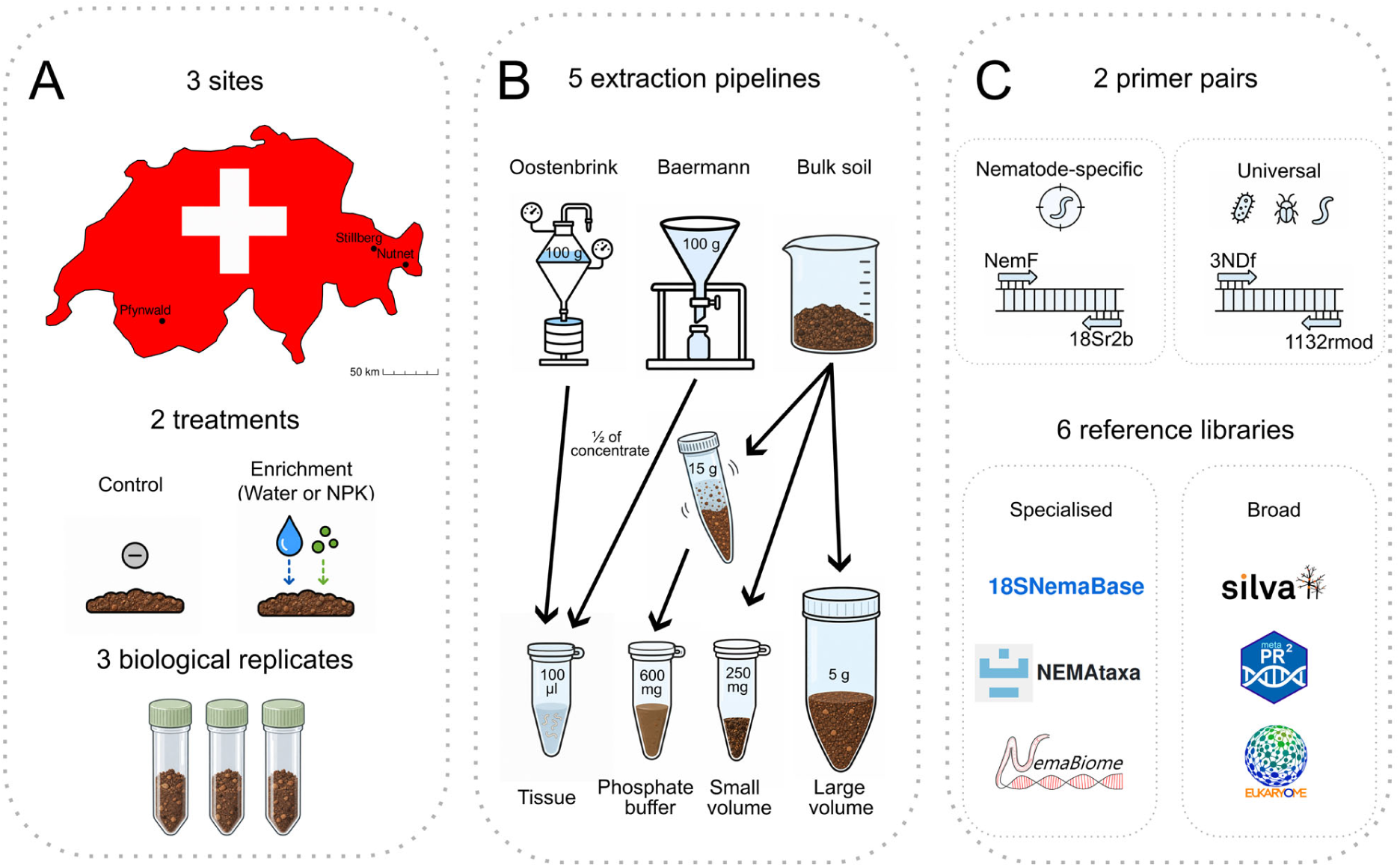
Experimental design. a) Sampling site locations within Switzerland and experimental site treatments. b) DNA extraction pipelines: Oostenbrink elutriator plus tissue extraction, Baermann funnel plus tissue extraction, extraction from bulk soil with Qiagen’s PowerSoil Pro kit (small volume), extraction from bulk soil with Qiagen PowerMax Soil kit (large volume) and extraction from bulk soil with a buffered slurry step followed by the Qiagen’s PowerSoil Pro kit. c) Primers used for the amplification of sequences and references libraries tested for the taxonomic assignment of sequences.

In the Stillberg experimental site, canton Grisons, 46°47ʹN, 9°52ʹE, 2100 m altitude, where an afforestation was carried out in 1975 above the natural treeline, the enriched *Pinus uncinata* plots received 30 kg N ha^−1^, 75% in the form of NH4^+^ and 25% as methylene urea every year from 2004 to 2016 (Möhl et al., 2019). In the Pfynwald experimental site, canton Valais. 46°18’N, 7°36’E, 615 m altitude, enriched *Pinus sylvestris* plots have received yearly since 2003 600 mm of water in addition of natural rainfall, via sprinklers (Bose et al., 2022). In the NutNet Switzerland alpine grassland experimental site, 46°63’N, 10°37’E, canton Grisons, 2320 m altitude, enriched experimental plots receive 10 g N m^-2^ as time-release urea, 10 g P m^-2^ as triple-super phosphate; 10 g K m^-2^ as potassium sulphate, applied yearly (Risch et al., 2020). In all cases, control plots had the same baseline conditions as enriched plots but they did not receive supplemental irrigation or the addition of fertilisers.

Three plots were sampled in each site from both the control and the enriched plot in the summer of 2023. Undisturbed soil cores (0 to 10 cm) were collected with a soil hammer, with 5 composite cores per plot manually mixed in a plastic bag. An aliquot from each sample was taken for direct DNA extraction and preserved at -20 °C. The rest of the soil was kept refrigerated at 4 °C, with extraction performed within 48 hours.

### Active and passive nematode extraction

Aliquots of 100 ± 0.5 g fresh soil were weighted for both the Baermann and the Oostenbrink pipelines. For the Baermann extraction, the soil was laid in a 15 cm diameter funnel on top of a sheet of 3-ply kitchen paper folded onto itself. The funnels were kept at room temperature and watered twice a day for four days (McSorley & Frederick, 2004). Watering was carried out from the top until the meniscus between the water surface and the kitchen paper covered 80% of the opening of the funnel. At the end of the 96 hours, the clamps at the bottom of the funnel were open to collect in a Falcon tube the bottommost 50 ml of slurry. This was then refrigerated at 4 °C for one hour and further reduced to 5 ml of concentrated slurry.

For the Oostenbrink elutriator, manufacturer’s instructions were largely followed. Specifically, the undercurrent was gradually moved from 60 to 40 l/h during the procedure until the layer with the nematode suspension could be diverted onto the outflow and through the superimposed 75 µm and 45 µm collection sieves. These were rinsed into a collection bowl, whose contents were then poured onto a concave watch glass laid at the centre of a 150 µm sieve on top of which a 2-ply kitchen paper towel and a milk filter were clamped. The clamped sieve was then carefully submerged into a collecting plate and let rest in the dark for 24 hours at 20 °C. The contents of the collecting plate were then poured into a jar. This was stored at 4 °C for an hour after which the supernatant was removed obtaining 5 ml of concentrated slurry.

### Moisture determination and counting

The concentrated slurry from both the Oostenbrink and the Baermann pipelines were poured onto a gridded Petri dish for counting under 20 X to 50 X stereoscopic magnification. After counting, the contents of the Petri dish were poured back into the tube, refrigerated overnight, concentrated to 2.5 ml, resuspended and divided into two aliquots, one for DNA extraction and the other kept in reserve. An additional 30 g fresh soil from each sample was heated at 105 °C for 24 hours for gravimetric moisture determination in order to convert nematode counts to dry soil estimates.

### DNA extraction

For the nematode slurries originating from the Oostenbrink and Baermann pipelines, the 1250 µl aliquot was centrifuged for 2 minutes at 13,000 *g* and the supernatant was removed leaving a concentrated 100 µl pellet (Figure 1b). 180 µl of Qiagen’s DNeasy Blood & Tissue ATL buffer and 20 µl of proteinase K from the same kit were added to the samples and incubated in a water bath at 56 °C for 4 hours (Resch et al., 2022). The extraction was then carried out with Qiagen’s DNeasy Blood & Tissue according to the manufacturer’s instructions.

For the small volume extraction, 250 (+-5 mg) of soil were inserted in bead lysis columns from Qiagen’s PowerSoil Pro kit and processed according to the manufacturer’s instructions, with a lysis step carried out on a MP Fastprep 24, 5 g bead beater at 5.5 m s^-1^ for two 30 s cycles separated by a 180 s cooling interval and a final elution in 80 µl.

The large volume extraction was carried out on 5 g (+-10 mg) of soil with a Qiagen PowerMax Soil kit according to the manufacturer’s instructions, with a 10 minute vortexing step.

The buffer extraction was carried out on 15 g of soil (+-10 mg). The soil was loaded in a 50 ml centrifuge tube, to which 15 ml of phosphate buffer (according to Taberlet et al., 2012) were added. The soil was shaken in an orbital shaker at 300 HZ for 30 minutes. 600 µl were loaded into the microcentrifuge tube, and processed with Qiagen’s PowerSoil Pro kit according to the manufacturer’s instructions.

### Amplification and sequencing

Amplification of 18S rRNA gene was done with two primer pairs: the nematode specific primer pair NemF/18Sr2b (Sapkota and Nicolaisen, 2015) referred hereafter as NEM primer and the universal eukaryotic primer pair 3NDf/1132rmod (Geisen et al., 2018), referred as UNI primer. PCR was carried out in a final 25 µl volume with reaction buffer 1x (KAPA HiFi buffer), 0.3 mM of KAPA dNTP mix, 0.3 µM of each primer, 0.5 U of KAPA HiFi Hotstart DNA polymerase (Roche, Switzerland), and 1-2 µl of template DNA. Negative controls with ultra-pure water were included in each PCR run to detect possible contaminations. The PCR temperature profile was: initial denaturation at 95 °C for 3 min, followed by 25 cycles of denaturation 98 °C for 20 s, annealing 67 / 63 °C for 15 s (NEM/UNI primer), and elongation 72 °C for 45 s, and a final elongation step at 72 °C for 5 min. Amplifications were performed in a thermal cycler (Veriti Pro, Applied Biosystems) and PCR products analysed by electrophoresis on 1.5% agarose TAE gels. Amplicons were then purified with the AMPure XP magnetic beads (Beckman Coulter, Inc., USA). Library preparation and sequencing was carried out as external service (Earlham Institute, UK), on a NextSeq 1000 P1 flow cell with 300 bp pair-end reads (Illumina). Alongside the samples, three extraction blanks were included and a custom-made mock community sample which included 10 nematode species.

### Bioinformatic processing and taxonomic assignment

Sequences were analysed using the DADA2 pipeline (Callahan et al., 2016) independently for each primer pair, with default parameters. For the universal primer pair 3NDf/1132rmod, only forward reads were selected due to non-overlapping reads. After filtering and de-noising steps 73% of the reads were retained, and 3,502 amplicon sequence variants (ASV) were identified with the Nem primer and 91% of the reads were retained and 36,283 ASV with the Uni primer. Taxonomic identification was performed by *dada2::assignTaxonomy* function, using six different reference databases, three universal eukaryotic, SILVA database v132 (Quast et al., 2013), PR2 v5.1 (Guillou et al., 2013) and EUKARYOME v1.9.4 (Tedersoo et al., 2024), and three nematode specific databases, NemaBase (Gattoni et al., 2023), NemaTaxa v1 (Baker et al., 2023) and Nemabiome v0.0.1 (Charrier et al., 2024).

Database was refined by three consecutive steps. 1) Lulu algorithm to reduce the number of erroneous ASVs and achieve more realistic biodiversity metrics (Frøslev et al., 2017), with a minimum match of 99.5% for the NEM primer and 99% for the UNI primer, and the rest parameters set as default. The election of these parameters was based on the results of the mock community sample. Lulu discarded 2,579 ASVs for the NEM primer and 3,797 ASVs for the UNI primer. 2) Minimal abundance filtering, by removing any reads that represent < 0.002% abundance, which discarded 739 ASVs and 4,688 reads for the NEM primer and 9,843 ASVs and 48,860 reads for the UNI primer. 3) Blank correction with “decontam” package (Davis et al., 2018), where contaminants were detected by prevalence with a threshold of 0.3. One ASV with 668 reads and three ASVs with a total of 23,883 reads were detected as contaminants for the NEM and UNI databases respectively. Final databases contained 3,502 ASVs and 23,258,259 reads for NEM primer, and 22,640 ASVs and 32,520,440 reads for UNI primer.

As reference libraries diverge wildly in taxonomic rank structure, making direct comparisons impossible, all assignments were mapped to the GBIF backbone taxonomy (GBIF Secretatiat, 2023). For each individual taxonomic assignment and each of the libraries, a recursive algorithm starting from the lowest assigned rank attempted a match with the *name_backbone* function of “rgbif” (Chamberlain et al., 2026) and moved upwards in case of no suitable hit. The mapping log from the algorithm was screened and manual corrections to account for bugs in the API interface were operated.

### Statistical analyses

For specimen counts, corrected for gravimetric moisture content, a linear model was fitted having extraction method, treatment and site as explanatory variables. Individual-based rarefaction curves of expected ASV richness were calculated using the *rarecurve* function in the R package “vegan” (version 2.7-2, Oksanen, 2024; Oksanen et al., 2008). Shannon diversity was analysed separately for all ASVs and nematode ASVs using linear models with extraction method, primer pair, their interaction, treatment, and site as predictors. Observed ASV richness was analysed with negative binomial regression in “MASS” (version 7.3-60.0.1, Ripley et al., 2026) using the same predictors plus log read depth. For each nematode order, its read count relative to other nematode reads was analysed with beta-binomial regression in “glmmTMB” (version 1.1.14, Brooks et al., 2026); likelihood-ratio tests assessed overall extraction-method and primer effects, and p-values were adjusted across orders and both effects using the Benjamini–Yekutieli method. The overall extraction-method effect in each primer-specific partial RDA was tested with 9,999 permutations under the reduced model using “vegan”, which also provided adjusted (R^2^).

Within each extraction method and primer pair, ASV counts were converted to within-sample relative abundances and Bray–Curtis dissimilarities were calculated. Site and binary enrichment status were assessed by sequential PERMANOVA with 999 unrestricted permutations, separately for all ASVs and nematode ASVs; reported p-values were unadjusted. For visualization, nematode ASV dissimilarities were subjected to principal coordinates analysis in “ape” (version 5.8-1, Paradis et al., 2024) separately for each method–primer combination.

All analyses were performed in R (version 4.3.3, R Core Team, 2025) through the RStudio integrated environment (version 2026.04.0, Posit team, 2024).

## Results

### Specimen recovery

Passive Oostenbrink extractions retrieved substantially more specimens than the active Baermann funnel method. Controlling for site and treatment the Oostenbrink pipeline yielded an estimated marginal mean count of 2316 specimens per 100 g of dry soil compared to Baermann’s 586 (*p* = 0.037). However, a higher variability in counts was observed in Oostenbrink extractions, with a pooled coefficient of variation of 1.498 compared to 0.656 for Baermann extractions (Figure 2).

**Figure 2:**
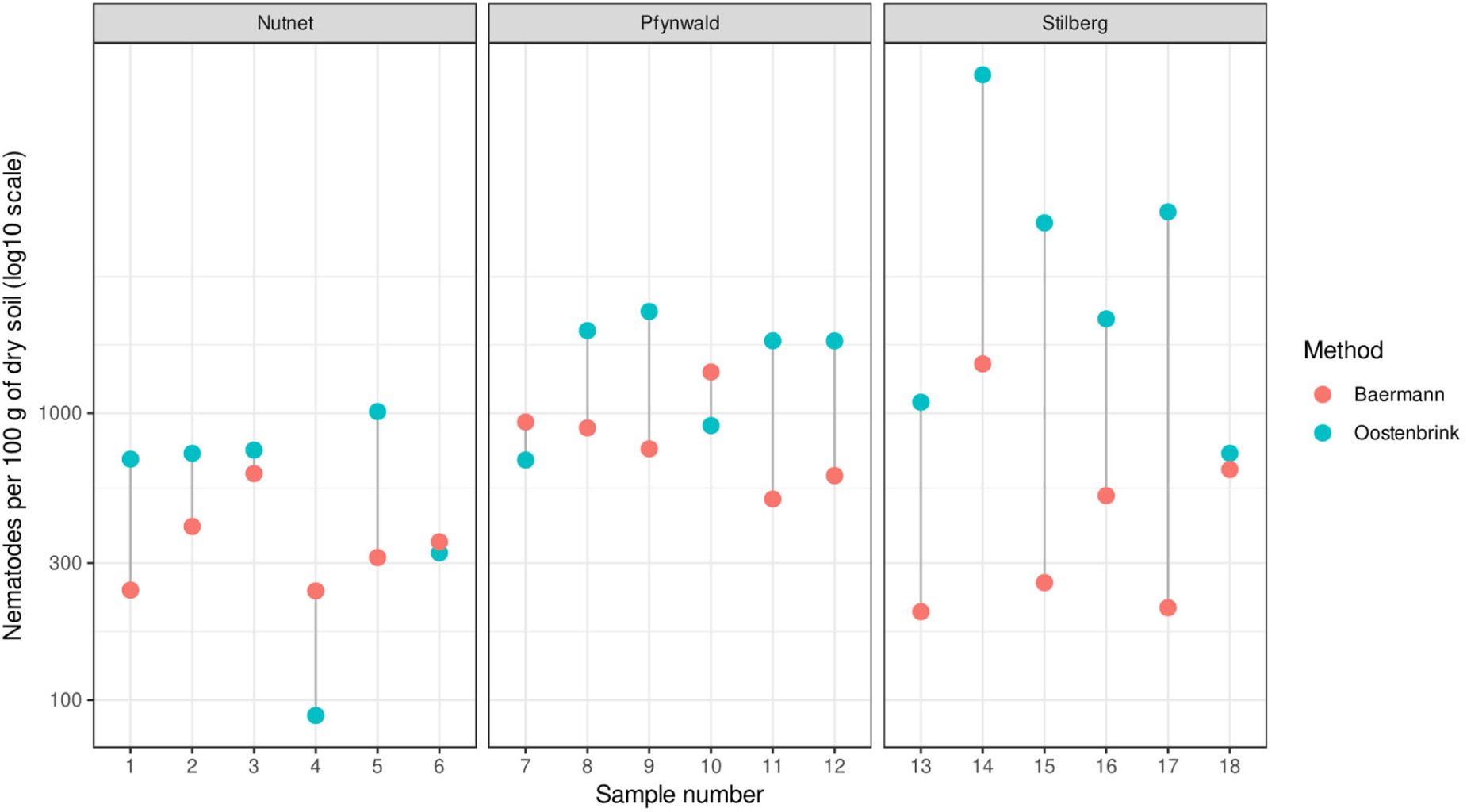
Counts of recovered nematodes per unit of dry soil with active (Baermann) and passive (Oostenbrink elutriator) extraction methods. The counts are based on an extraction of 100 g of fresh soil and converted according to gravimetric moisture content.

### ASV diversity and richness

Considering all ASVs, the Oostenbrink pipeline resulted in a higher Shannon’s diversity score (+0.441, 0=0.005, Figure 3a) compared to the Baermann pipeline, and so did the small volume extraction (+2.180, p<0.001), the large volume extraction (+2.228, p<0.001) and the buffered extraction (+1.749, p<0.001). The NEM primer pair resulted in a lower Shannon’s diversity measure (-0,515, p=0.001). Significant negative interaction effects were also registered for NEM combined with the three bulk soil extraction methods. For ASVs that were assigned as nematodes according to the Eukaryome library, a similar pattern was observed. Specifically, compared to the Baermann pipeline baseline, all methods resulted in higher scores (Oostenbrink +0.561, p<0.001; small volume +2.351, p<0.001); large volume +2.362, p<0.001; buffered +1.950, p<0.001). No significant individual effect was detected for the NEM primer pair, but all three bulk soil extraction methods had a negative interaction effect when combined with NEM.

**Figure 3:**
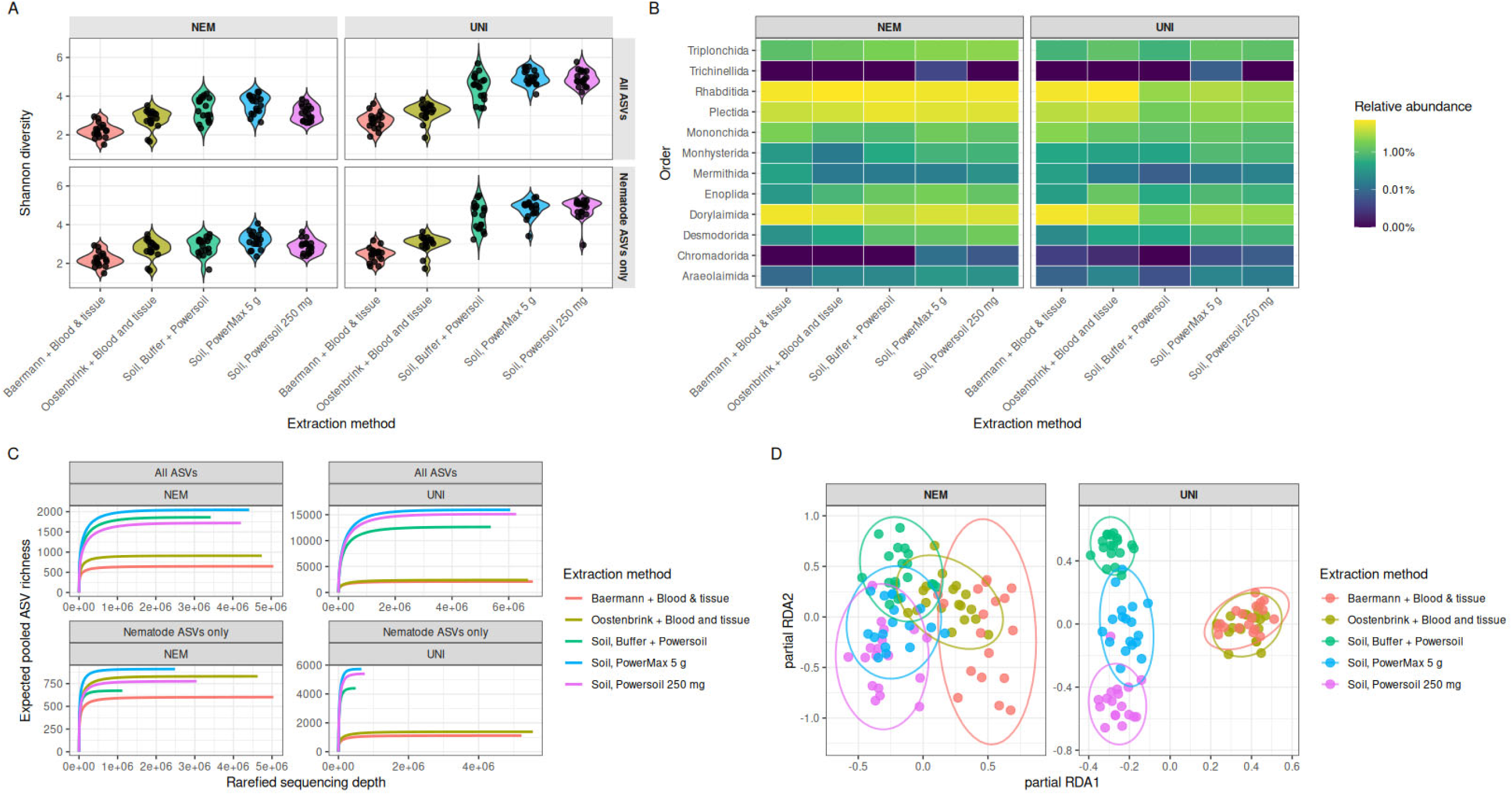
a) Violin plot of Shannon’s diversity indices calculated for ASV distributions obtained from different pipelines. Each point represents a single sample. b) Relative abundance heatmap for the 12 nematode orders represented in the sequenced samples. c) Rarefaction curves for pooled samples grouped for each extraction method and primer pair. d) Partial RDA performed separately for each primer on Hellinger-transformed nematode ASV abundances, with extraction method as the explanatory variable and site and treatment as conditioning variables.

Total ASV richness is also computed along these lines (Figure 3c). The lowest ASV number was registered in the Baermann pipeline, with the Oostenbrink pipeline (+0.313 log richness, p=0.001), and the small volume (+2.443, p<0.001), large volume (+2.404, p<0.001) and buffered extraction (+2.224 p<0.001) all resulting in significantly higher ASV counts. The NEM primer pair resulted in a 0.445 log richness decrease (p<0.001) and all three bulk soil extractions had significant negative interactions with the NEM primer. Focusing only on sequences identified as belonging to nematodes by the Eukaryome target references, we can measure positive effect on richness compared to the Baermann pipeline for the Oostenbrink pipeline (+ 0.313, p=0.001), the small volume extraction (+2.443, p<0.001), the large volume extraction (+2.404, p<0.001) and the buffered extraction (2.224, p<0.001). Again, the NEM primer pair induced a significant decrease in observed ASV richness (-0.445, p<0.001) alone and in its interaction with all three bulk soil extraction methods.

### Extraction-specific taxonomic biases

Several major nematode orders showed biases in detection and quantification attributable to either primer selection or extraction methods (Figure 3b). In particular, *Mononchida* and *Monhysterida*, both primer choice and extraction method resulted in significantly different relative abundances (p<0.001). For *Desmodorida*, *Dorylaimida*, *Enoplida*, *Triplonchida* (p<0.001) and *Plectida* (p<0.01) only the extraction method proved to be a significant driver of differential abundance.

For both primers a statistically detectable association between extraction method and nematode ASV composition is registered, after accounting for trial and the binary treatment (p<0.0001). However, the variability explained by extraction itself is quite limited (adjusted R^2^ of 0.100 for the UNI primer and 0.055 for the NEM primer). The tissue-based methods tend to cluster closer together, in particular when coupled with the UNI primers (Figure 3d).

### Community analysis and experimental signal

All five extraction methods recovered a site-associated difference in community composition with both primer pairs, for all ASVs and for nematode ASVs alone (PERMANOVA R^2^ between 0.331 and 0.464, unadjusted permutation, p=0.001 in all 20 analyses, Figure 4a). In contrast, the enrichment contrast was detected only by the UNI primer set coupled with the small volume extraction (unadjusted p=0.040). The visual assessment of treatment and environmental clustering in separate bidimensional ordinations shows clustering patterns that largely preserve the same centroid distances and distribution across methods and primers, with a less marked site signal resulting in overlapping ellipsoids in the Baermann pipeline (Figure 4d).

**Figure 4:**
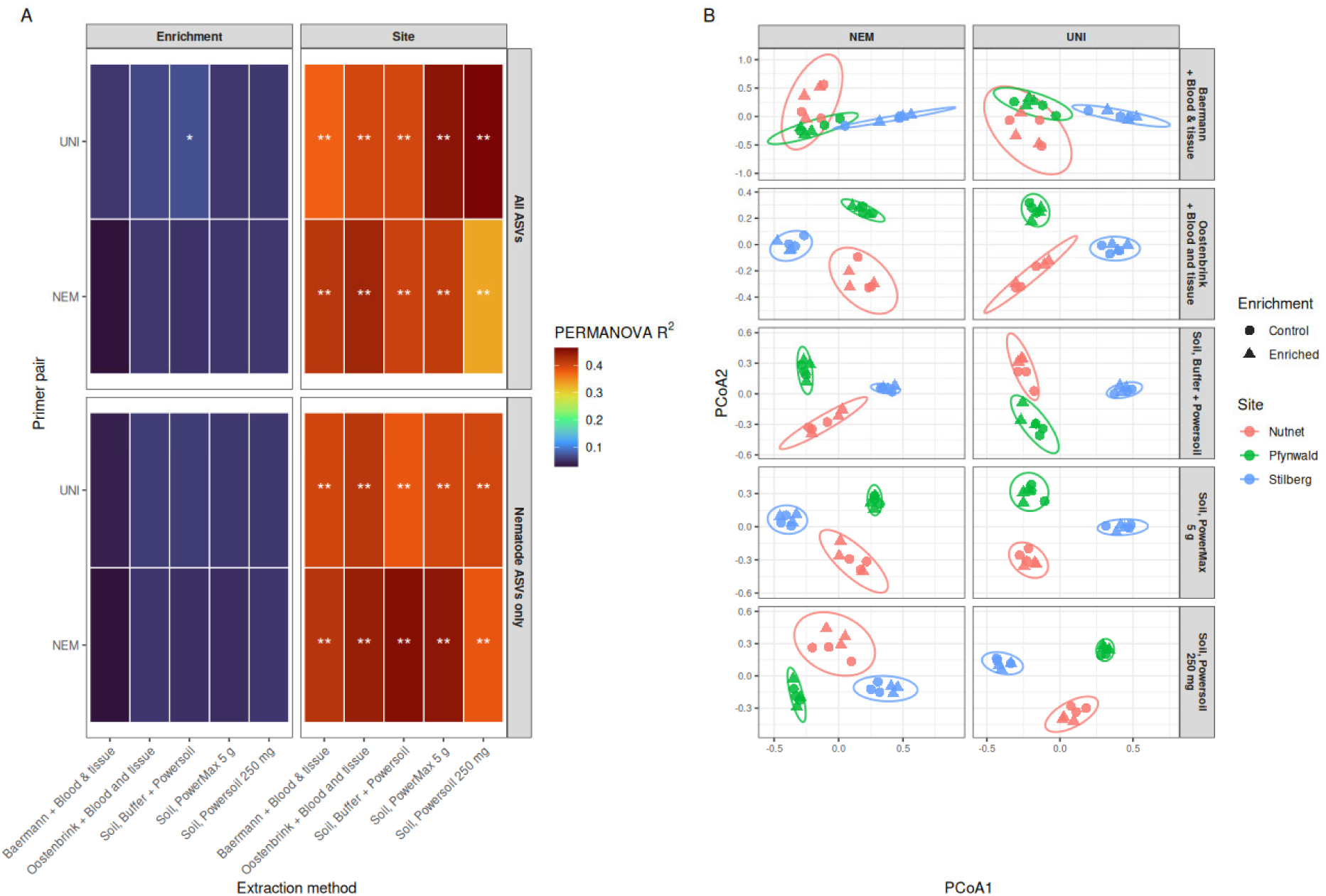
a) PERMANOVA (R^2^) for site and enrichment, calculated separately for each extraction method, primer pair, and ASV subset. Asterisks indicate unadjusted permutation p-values. b) Principal coordinates analysis (PCoA) of nematode ASV communities. Each point represents a sample; colours indicate site, shapes indicate enrichment status, and concentration ellipsoids of the 0.95 t-distribution are drawn for each site. PCoA was calculated separately within each method–primer panel, so axis positions are not comparable between panels.

### Library resolution and agreement

Reference libraries differed substantially in the proportion of sequences assigned to Nematoda (Figure 5a). With NEM primers, nematode assignments represented 9.9–100% of reads and 6.8–100% of ASVs across libraries; with UNI primers, the ranges were 33.8–100% of reads and 5.1–100% of ASVs. When using universal eukaryotic reference databases, results where similar among the three databases, with 70.3-74.5% of reads assigned to Nematode and 38.1-42% of the taxa when amplified with the NEM primer, and 33.7-35.8% of the reads and only 5.1-6.6% of the taxa when amplification was done with the UNI primer. Regarding nematode-specific databases, Nemabiome assigned all retained reads and ASVs to Nematoda, while NemaBase assigned 81.9% of NEM reads and 93.5% of UNI reads to Nematoda. NemaTaxa assigned 9.9% of NEM reads to Nematoda and produced no hits with UNI.

**Figure 5:**
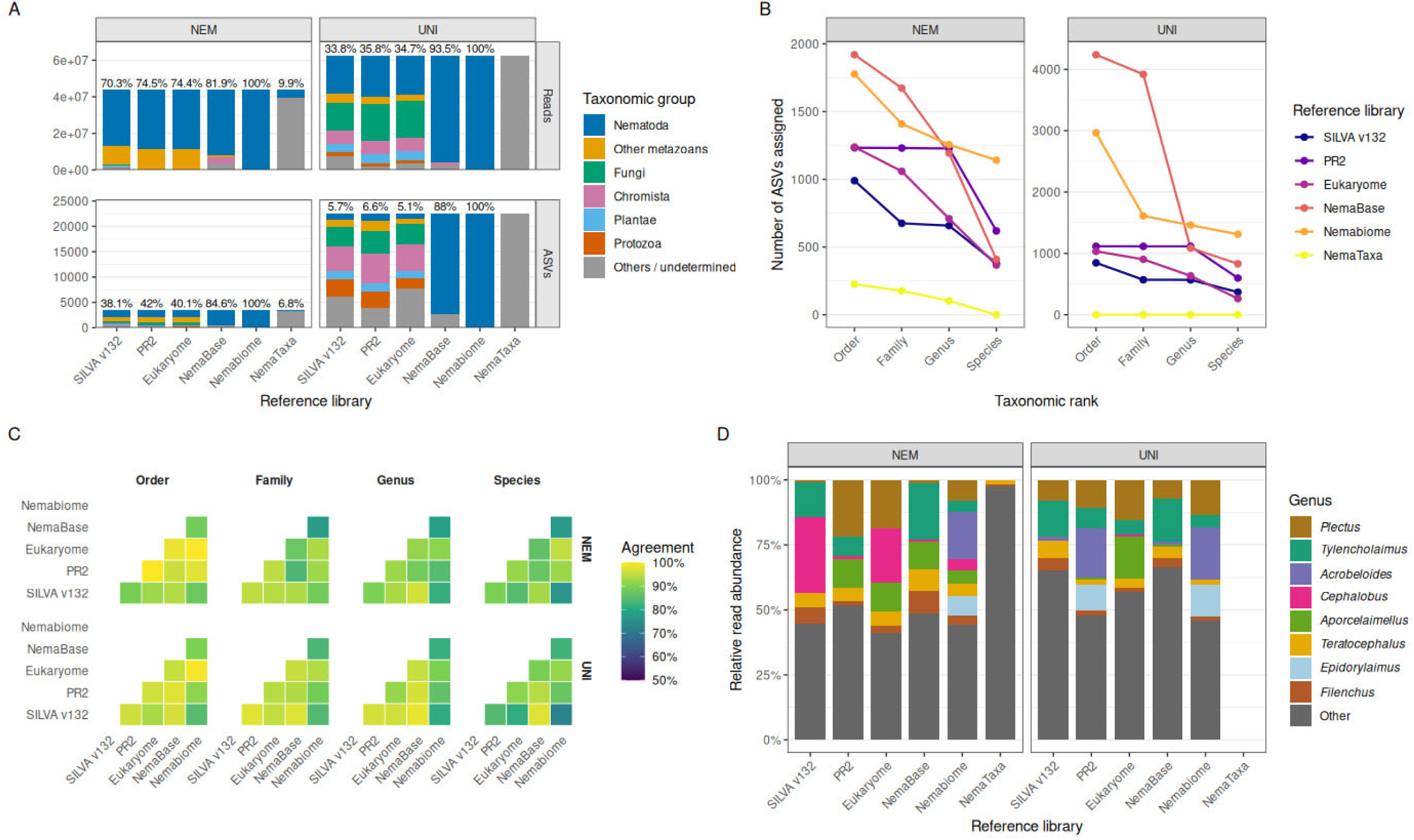
a) Proportion of reads and ASVs per reference library that were assigned to different clades according to the GBIF mapping of the named taxon. b) Number of ASVs assigned by the reference libraries at different taxonomic ranks. c) Agreement heatmap of taxonomic assignment based on taxa that both libraries in a comparison recognise as nematodes. d) Proportion of reads assigned to the 8 most abundant nematode genera across the pooled samples, divided by library.

The number of nematode ASVs with finer-rank assignments also varied (Figure 5b). Nemabiome provided the most species-level assignments for both NEM (1,141 ASVs) and UNI (1,313 ASVs). NemaBase provided the most order-level assignments (1,920 and 4,239 ASVs, respectively), although its genus-level counts were lower (1,193 and 1,087). Among ASVs classified as nematodes and assigned at the same rank by both libraries, pairwise agreement was generally high: order-level agreement ranged from 87.7–100% for NEM and 84.8–99.4% for UNI (Figure 5c). Species-level agreement was more variable, ranging from 76.0–95.0% and 72.1–97.5%, respectively. The relative read profiles of assigned nematode genera also changed across libraries (Figure 5d); for example, *Cephalobus* comprised 29.3% of genus-assigned NEM reads with SILVA v132 but 0.9% with PR2, whereas *Plectus* comprised 0.8% and 21.7%, respectively.

## Discussion

The Oostenbrink passive elutriator recovered substantially higher numbers of nematodes from soil compared to the low-cost and high-throughput Baermann funnels, at the expense of a higher initial investment, maintenance and time required per sample. The numeric advantage extended down the pipeline, yielding significantly higher ASV richness and diversity. The higher recovery efficiency is consistent with existing literature (Filgueiras et al., 2025; Thiruchchelvan et al., 2024; Verschoor & Goede, 2000). However, the higher yields came with a higher mean-controlled variability, whereas Baermann funnels were remarkably more consistent.

The two pipelines involving extraction of nematodes from soil scored slightly lower than the bulk soil methods in terms of Shannon’s diversity compared to the bulk soil ones, irrespective of the primer used, which is in disagreement with previous studies (Donhauser et al., 2023). In terms of ASV richness, the differences were considerable. The largest gap between rarefaction curves was observed when using the UNI primer, which was to be expected. Separating the nematodes from the matrix reduced by a great amount the number of non-nematode sequences in the sample, which is why the technique is still widely adopted in large scale studies when nematodes are the primary focus of the study (Akanwari et al., 2026; Donhauser et al., 2023; Kawanobe et al., 2021). With NEM specific primers, the difference was drastically reduced, even more so when only nematode sequences are taken into account, indicating the good specificity of separation pipelines. While at order level, in a reference-standardised analysis using Eukaryome assignments, the different methods largely agreed in identifying the most abundant taxa, significant effects of primer and even more so extraction method, were detected. Once again, the clustering tended to separate the direct extraction from soil method from the Oostenbrink and Baermann pipelines, which is particularly evident with the UNI primer set. Reasons for these differences can be identified in annealing conflicts and suppression of rare taxa when non-target sequences are dominant (Akanwari et al., 2026; Ficetola et al., 2024; Gold et al., 2023; Sikder et al., 2020) and lysing bias in nematode pellets from separated tissue extraction (Rybarczyk-Mydłowska et al., 2026; Waeyenberge et al., 2019), with all effects compounded by primer-specific biases (Akanwari et al., 2026; Sapkota & Nicolaisen, 2015; Sikder et al., 2020). In contrast with some of the literature on the subject (Dopheide et al., 2019; Kageyama & Toju, 2022; Wang et al., 2026) the quantity of soil used in the extraction did not affect dramatically richness, diversity and community structure, the three bulk soil pipelines being largely aligned throughout.

As for community level ordination-type analysis and the detection of environmental gradients, the prospects for agreement among different pipelines are much more robust. All methods provided a largely similar clustering pattern and comparable centroid distances using dimension reduction on dissimilarity matrices, and all methods were largely aligned in detecting and quantifying site and treatment specific signals. This is in complete agreement with a large and growing body of literature according to which beta-diversity patterns and ordination structures tend to maintain higher congruence in the face of widely different methodological choices (Gonzalez-Saldias et al., 2026; Hajibabaei et al., 2019; Van den Bulcke et al., 2023).

The taxonomic-reference naive or standardised measures discussed up to now appear to be relatively less important in magnitude than the divergence observed while adding different taxonomic reference libraries to the analysis. The two main clusters are predictably represented by generalist or pan-eukaryotic libraries and nematode-specific references. Within this second cluster, the poor performance of the NemaTaxa assignment should not be read as a quality judgment about this highly-refined and skilfully-curated library but rather as a warning about coupling it with different sets of primers as the ones used to design this database. As for the other two nematode-specific references compared, NemaBase and Nemabiome, it has to be noted that, using default parameters in the algorithm, they tend to assign to nematode taxa ASVs that broad-scope libraries confidently assign to other taxa. Careful tuning of bootstrapping and confidence threshold should be adopted to reduce the risk of overassignment. The problem is particularly acute when such a library is used in combination with a generalist primer set, where the assignment of over half of the reads pinpointed to other clades by broad libraries gets mapped to nematode taxa. The risk of false positives is known to many metabarcoding libraries (Chorlton, 2024; Murali et al., 2018) and has been detected for nematodes too (Macheriotou et al., 2019).

A problem conversely affecting broader-targeted taxonomic libraries more severely than specialised ones is the rapid assignment decay along the rank hierarchy. Confident assignment of a broad range of eukaryotic taxa to large groups and reduction of false positives seems to be pitched in a trade-off with hits at genus or species level, which is widely reported in literature comparing specialised and broad libraries (deWaard et al., 2019; Guillou et al., 2013; Mugnai et al., 2023). This effect, when interpreted through the lens of another phenomenon emerging in our study, namely the high rate of paired cross-reference library agreement in cases where a sequence is assigned to the same target clade (i.e. Nematodes), introduces the possibility of combining broad and specialised libraries together, to reduce the rate of false positives while at the same time preserving the resolution at the lower ranks. Nevertheless, the poor rate of vertical alignment and the idiosyncratic treatment of ranks in Linnean derived taxonomy across different references, makes the operation a complex undertake (De Santiago et al., 2025). In addition, even within the realm of specialised libraries with a narrow taxonomic focus and more thorough alignment at higher ranks, the differences in detected community make-up at genus level or lower still require significant harmonisation.

## Conclusions

Our study clearly demonstrates that the choices of methods for nematode biodiversity studies using molecular tools can have a big impact on the results. These methods include the extraction step, following with the primer pair for amplification and finishing with the reference taxonomic database for taxonomic identification. Observed differences were more pronounced if comparing the nematode extraction *vs* soil extractions. In particular, direct soil extractions co-extracted many non-nematode taxa, especially if amplified with a universal primer, which can significantly reduce the accuracy of nematode community indexes and other nematode-based metrics. The choice of the reference database strongly affected the taxonomic identification, which calls for extreme caution when making this selection. However, the ecological signal of the obtained nematode community structure was consistent across methods, highlighting beta-diversity metrics as robust against methodological choices.

## Notes

### Competing Interest Statement

The authors have declared no competing interest.

